# Melanin pathway genes regulate color and morphology of butterfly wing scales

**DOI:** 10.1101/234831

**Authors:** Yuji Matsuoka, Antónia Monteiro

**Affiliations:** Department of Biological Sciences, National University of Singapore, 14 Sciences Drive 4, 117543 Singapore; Science Division, Yale-NUS College, 10 College Avenue West, 138609 Singapore

## Abstract

The cuticular skeleton of a butterfly wing scale cell serves both as a substrate for the deposition of pigments and as an exquisitely finely-sculpted material responsible for the production of structural colors. While cuticle rigidity and pigmentation depend to a large extent on the end products of a branched biochemical pathway – the melanin pathway – little is known whether genes in this pathway also play a role in the development of specific scale morphologies that might aid in the development of structural colors. Here we first show that male and female *Bicyclus anynana* butterflies display differences in scale size and scale morphology but no differences in scale color. Then we use CRISPR/Cas9 to show that knockout mutations in five genes that function in the melanin pathway, *TH, DDC, yellow, ebony,* and *aaNAT,* affect both the fine structure and the coloration of the wing scales. Most dramatically, mutations in *yellow* led to extra horizontal cuticular laminae on the surface of scales, whereas mutations in *DDC* led to taller and sheet-like vertical cuticular laminae throughout each scale. We identify some of the first genes affecting the development of scale morphology, and whose regulation and pleiotropic effects may be important in creating, as well as limiting, the diversity of structural as well as pigmentary colors observed in butterflies.

## Introduction

The exoskeleton of insects not only provides important structural support, but is also used as a canvas for the deposition of pigments or for the sculpting of structural colors, used in camouflage and mate signaling. The skeleton is made up of cuticle, an extracellular matrix comprised of chitin fibers, cuticular proteins, lipids and pigments (Moussian, 2010). One of the most common pigments found in insect cuticle is melanin, and some melanin pathway products also take part in cuticle hardening/sclerotization, making cuticular melanization in insects tightly linked with cuticular sclerotization. For example, the cross-linking of melanin pathway molecules with nucleophilic amino acid residues and cuticular proteins confers cuticule stiffness (Xu et al., 1997, Kerwin et al., 1997, Andersen, 2005, Suderman et al., 2006). The interaction among all these cuticle components is, thus, important in the assembly of the insect exoskeleton and in its coloration (Xiong et al., 2017).

Given the interdependency of coloring and hardening cuticle we sought to explore whether mutations in the enzymes that control the flux of chemical precursors across pigment biosynthetic pathways affect both the spatial patterns of cuticle deposition in addition to its coloration.

We chose to examine the effects of pigment enzyme mutations on both the color and intricate morphology of butterfly wing scales because scales of different color within a wing often display different cuticular micro-morphologies (Ghiradella et al., 1972, Vukusic et al., 2000, Janssen et al. 2001, Stavenga et al., 2004, Siddique et al., 2015). This suggests that the two processes might be genetically linked. Furthermore, scale pigmentation and morphological patterning coincide during pupal wing development. For instance, melanin pigments are deposited at late development stages, with a subtle time lag between the deposition of yellows, browns, and black melanin pigments (Koch 1998, Wittkopp and Beldade, 2009), and this process correlates with the development of fine morphological features of scales such as the development of the longitudinal ridges and crossribs (Waku and Kitagawa, 1986; Ghiradella, 1989) (Fig. 1). In addition, most melanin pigmentation enzymes are expressed at these same late stages (Nishikawa 2013, Connahs et al. 2017, Zhang et al. 2017), indicating that they might play a dual role in scale construction and pigmentation. We hypothesized, thus, that molecules that determine color in butterfly wings might also regulate scale morphology.

**Figure 1.**
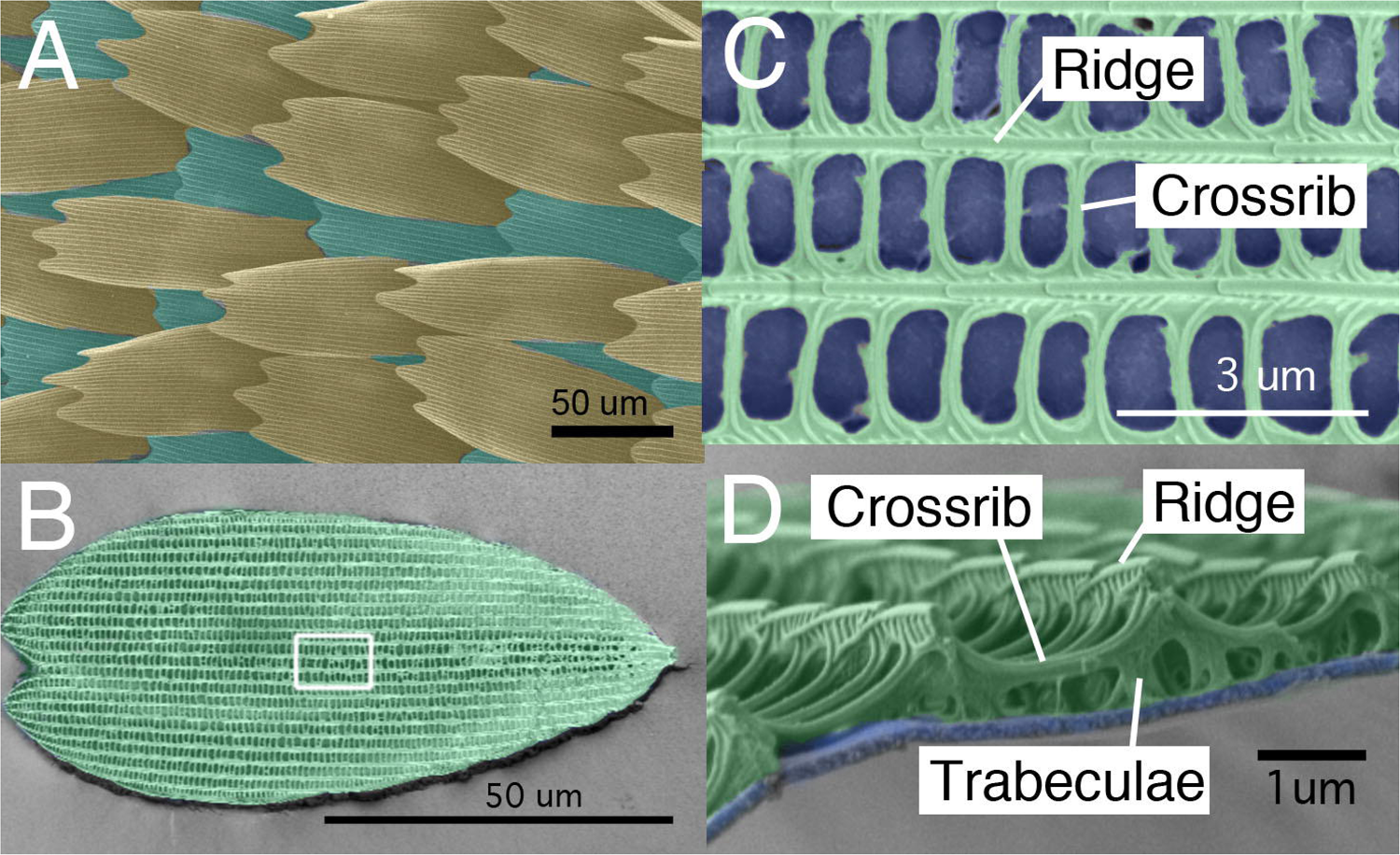
Wing scale arrangement and scale structure of *B. anynana*. (A) Arrangement of butterfly wing scales with cover (tan) and ground (green) scales alternating each other along vertical rows. (B) Individual ground scale where the white square indicates the region magnified in C. (C) Ridges run along the longitudinal axis of the scale, and crossribs connect neighboring ridges, resulting in the delineation of open spaces, or windows (blue). (D) Cross-sectional view of a similar area as in C, showing a smooth lower lamina (blue) and a more intricately patterned upper lamina (green). The upper lamina connects to the lower lamina via the trabeculae. Scales and features of scales are artificially colored.

We focused on enzymes from two main pigmentation pathways, the melanin and the ommochrome synthesis pathway. The melanin biosynthetic pathway comprises a well-studied branched series of chemical reactions that can produce five different final molecules: two types of eumelanin, dopa-melanin and dopamine-melanin, pheomelanine, *N*- *β* -alanyldopamine (NBAD), and *N*-acetyldopamine (NADA) (Fig.2a)(Galván et al., 2015 and Arakane et al., 2009). Eumelanins are made from dihydroxyphenylalanine (DOPA) or from dopamine through additional reactions catalyzed by the *laccase2* and *yellow* family gene products to produce black (dopa-melanine) and brown pigments (dopamine-melanin) (Andersen, 2005). Recently, pheomelanine (a reddish-yellow pigment) was identified as an additional final product of the melanin pathway in insects (Speiser et al., 2014, Galván et al., 2015, Jorge Garcia et al., 2016, Polidori et al., 2017) (Fig. 2a). NBAD and NADA are sclerotizing precursor molecules having a yellow color, and being colorless, respectively. These molecules are made from dopamine through a reaction catalyzed by the *ebony* gene product and the *aaNAT* gene product, respectively. The ommochrome pathway typically produces yellow, orange and red pigments (Reed et al., 2008, Ferguson and Jiggins, 2009). The *vermillion* gene product, tryptophan oxidase, is the most upstream enzyme in the pathway, whereas *white* and *scarlet* code for a transporter protein which allow ommochorme pigment precursors to be incorporated into pigment granules (Mackenzie et al., 2000) (Fig. 2b).

**Figure 2.**
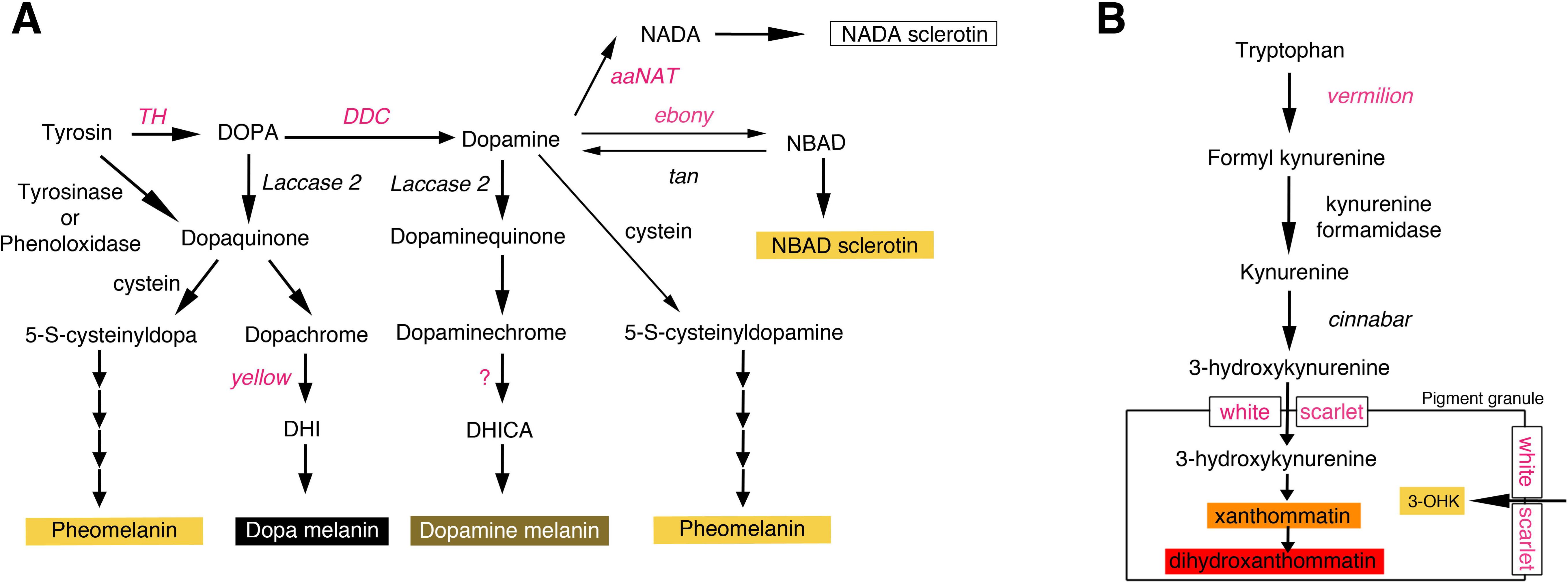
Proposed melanin and ommochrome biosynthesis pathway in insects. (A) The melanin biosynthesis pathway produces up to five different molecules: pheomelanin (reddish-yellow pigment), dopa-melanin (black pigment), dopamine-melanin (brown pigment), NADA sclerotin (colorless molecule) and NBAD sclerotin (yellow molecule). NADA and NBAD sclerotin contribute to the sclerotization of cuticle when crosslinked with cuticular proteins. Dopa-melanin and Dopamine-melanin are thought to be generated by the oxidative polymerization of 1-carboxyl 5, 6-dihydroxyindole-2-carboxylic acid (DHI) and 5,6-dihydroxyindole (DHICA), respectively. Pheomelanin is generated by the oxidative polymerization of cysteinyldopa or cysteinyldopamine. The scheme is modified from Galván et al. (2015) and Arakane et al., (2009). (B) The ommochrome pathway produces up to three different pigments: 3-hydroxykynurenine (3-OHK) (yellow), xanthommatin (orange), and dihydroxanthommatin (red). (The scheme is modified from Reed et al., 2008). Highlighted in red are the enzymes investigated in this study.

We chose to examine the functions of five melanin and three ommochrome biosynthesis genes in controlling the color and morphology of the wing scales of *Bicyclus anynana*, an African butterfly (Lepidoptera; Nymphalidae; Satyrinae), whose brown, beige, black, and gold colors likely result from pigments in these pathways (Fig. 2). We first examined differences in scale color and morphology across the sexes. Then we used CRISPR/Cas9 to knock-out these eight genes. Recent CRISPR/Cas9 mutagenesis in *B. anynana* as well as in other butterfly species targeted a few melanin genes and reported color disruptions in all species (Zhang et al. 2017). Here, in addition to the previously described color disruptions in *yellow* and *ebony* in *B. anynana* (Zhang et al. 2017), we targeted the function of three other melanin biosynthesis pathway genes, *Tyrosine Hydroxylase (TH), DOPA decarboxylase* (*DDC*), and *Arylalkylamine N-Acetyltransferase* (*aaNAT)* and three ommochrome biosynthesis pathway genes, *vermilion, white*, and *scarlet,* with the aim of identifying the enzymes and biosynthesis products needed for generating each of the scale colors in *B. anynana.* Finally, we examined, for the first time, whether some of these enzymes/molecules contribute to both color and morphology of scales.

## Results

### Morphology, but not color, differs between male and female scales

Before conducting analysis on pigment biosynthesis gene mutants, we first analyzed the differences between wild type male and female scales. Scales of *B. anynana* represent typical butterfly scales, containing a smooth lower lamina and a more intricately patterned upper lamina with longitudinal ridges that run along the proximal-distal (P-D) axis, and connect to the lower lamina via short pillars called trabeculae (Ghiraldela, 1976, Wasik et al. 2014) (Fig.1). Each longitudinal ridge is connected via thinner crossribs, resulting in the delineation of open spaces, or windows (Fig. 1). Butterflies also have two types of scales, cover and ground, that alternate along a row, and which can differ in both structure and pigmentation. We found that brown and beige scales have much larger window sizes than eyespot-forming scales (white, black, and gold) (Suppl. Fig.1). We then examined each of these scale types independently in each sex. We found a variety of differences between the sexes in scale size and morphology but no difference in color (Suppl. File 1). The trend was similar in ground scales (Suppl. File 2). Based on these results, we decided to focus exclusively on males for the subsequent study of the effects of mutations of melanin and ommochrome pathway genes on color and scale morphology.

### Targeted mutagenesis against melanin and ommochrome biosynthesis genes

Using CRIPR/Cas9 we disrupted the functions of five melanin biosynthesis pathway genes, *TH, DDC, yellow, ebony,* and *aaNAT,* and three ommochrome pathway genes, *vermilion, white,* and *scarlet* (injection details, numbers, etc., are summarized in Suppl. Table 1). We scored phenotypes in G_0_ mosaic individuals.

#### *Tyrosine Hydroxylase (TH)* mutants

TH catalyzes tyrosine into DOPA, which is the initial step of the melanin pathway (Fig. 2). Removing TH function is known to eliminate all the melanin pathway products in a variety of arthropods including in *B. anynana* (True et al., 1999, Liu et al., 2010, Liu et al., 2014, Zhang et al. 2017; Fig. 2). Three out of 21 larvae showed lack of black pigment on the head capsule, which is usually black in first and second instars (Suppl. File3a). Three out of six butterflies successfully emerged from pupae. Wing scales of all colors were missing in some of the *TH* mutant clones (Fig. 3a). In other clones (perhaps heterozygous clones with a single copy of *TH* mutated), scales were present but had less pigment and were curled. White scales, even in putatively heterozygous clones contiguous with other colored regions, were affected in more extreme ways and were always absent. Together, these results indicate that *TH* is required for larval head pigmentation, and for scale overall development, pigmentation and structural rigidity. These phenotypes are similar to those observed in other butterfly species (Zhang et al. 2017).

**Figure 3.**
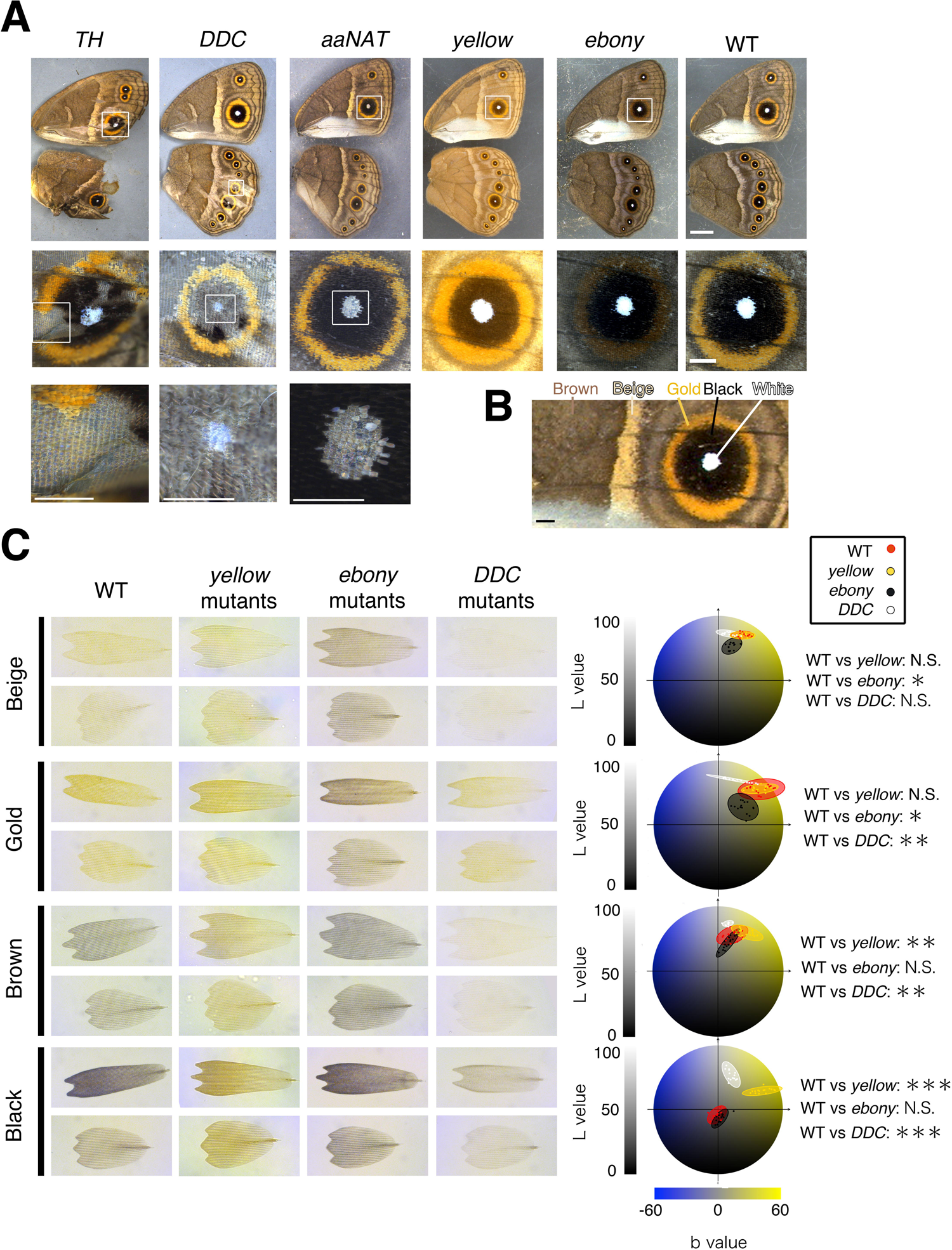
Melanin biosynthesis gene mutants in *B. anynana*. (A) Representative pictures of each melanin gene mutant. *TH:* Scales of all colors failed to develop in mutant *TH* clonal tissue. *DDC*: Scale development was also disrupted but not as frequently as in *TH* mutants. White scale development was always disrupted in mutant clones but gold, brown, and black scales became paler and curled in presumed heterozygous mutant clones. In *aaNAT* mutants white scale development alone was disrupted. In *yellow* mutants brown and black scales became lighter and in *ebony* mutants beige and gold scales became darker. Scale bar indicates 5 mm in low magnification images (top row), and 1 mm in higher magnification images (middle and bottom rows). (B) Colored areas that were sampled for individual scale measurements. Scale bar indicates 1 mm. (C) Transmission photos of individual mutant and WT cover and ground scales. Right diagrams show the color space occupied by cover scales on the L*a*b color space. Each colored ellipse represents measurements from 5 scales from 3 different male individuals (N=3 for statistical analyses).

#### *DOPA decarboxylase (DDC)* mutants

DDC catalyzes the production of dopamine from DOPA (Fig. 2). By disrupting DDC’s function, butterflies are expected to be able to produce only dopa-melanin (black pigment), however, they might still be able to produce pheomelanin if that part of the pathway is active in *B. anynana* wing scale cells. Five out of 42 larvae lacked black pigment on their heads (Suppl. File3a). From two of the seven butterflies that emerged, the black eyespot scales became gray, the brown and beige scales became whitish, the gold ring region became paler while still retaining a gold color, and the white eyespot center scales were absent (Fig. 3a). Furthermore, all scales were mostly curled. All these phenotypes were similar to the mild *TH* mutant phenotypes. L*a*b color space quantitative analyses (Hirschler, 2010) supported the visible color changes described: *DDC* mutant scales, other than beige scales, were significantly lighter (higher L value) and less yellowish in color (lower b value) compared to WT scales (Fig.3b). Our results indicate that dopamine is an important precursor for the pigments contained in most wing scales and also a required molecule in the development of white scales. The persistence of black and gold pigments in the black and gold mutated scales, respectively, indicate that dopa-melanin is likely present in the black scales, and other non dopamine-derived pigments (possibly pheomelanin or an ommochrome pigment) is present in the gold scales. These results also indicate that *DDC*, similarly to *TH*, is required for scale hardening across all the colored scales.

#### *Arylalkylamine N-Acetyltransferase (aaNAT)* mutants

AaNAT catalyzes the conversion of dopamine to NADA sclerotin (colorless) (Fig.2). Wild-type *B. anynana* naturally change their head capsule color from black in the third instar to light brown in the fourth instar (Suppl. File3b). From 20 larvae showing a normal black head as third instars, 16 retained their black head capsule color till the last larval instar (Suppl. File3b) This phenotype is similar to the phenotype observed in *Bombyx aaNAT* RNAi larvae (Long et al., 2015). Four out of 20 butterflies lacked white scales in multiple eyespot centers (Fig.3a), but no other scale abnormalities were detected. These results indicate that *aaNAT* is required for brown head capsule pigmentation in late larvae, the lighter color achieved presumably by shunting DOPA away from dopa-melanin production and into NADA sclerotin production. *aaNAT* is also required for the development of the white, structurally colored scales of the eyespot centers.

#### *yellow* and *ebony* mutants

Color changes in both *yellow* and *ebony* mutants in *B. anynana* were previously broadly described (Zhang et al. 2017), but here we quantified those changes using the L*a*b color space analysis (Fig.3b). In *yellow* mutants, the darker colored scales (black and brown) became significantly lighter relative to WT scales whereas the lighter colored scales (gold, beige, and white) did not change in color. This indicates that *yellow* is active only in darker colored scales. We infer that in these scales *yellow* is acting to convert DOPA to dopa-melanin.

In *ebony* mutants, the result was opposite to that of *yellow* mutants, that the lighter colored scales (gold and beige) became significantly darker relative to WT scale whereas the darker colored scales (black and brown) did not change in color (Fig. 3b) This indicates that ebony is active only in the lighter colored scales to convert dopa-melanin to NBAD sclerotin (yellow). The removal of *ebony* makes these scales less yellow (lower b) and darker (lower L) (Fig. 3b).

#### *Ommochrome* biosynthesis pathway likely does not contribute to wing coloration in *B.anynana*

To examine whether the ommochrome pathway contributes to the production of the gold color in *B. anynana*, we disrupted the function of three ommochrome biosynthesis pathway genes, *vermilion, white,* and *scarlet.* We confirmed the presence of indel mutations in groups of embryos after injection of gRNA targeting each of these genes along with Cas9, but did not observe any wing phenotypes in adults (Suppl. Fig4). The *white* mosaic mutants, however, showed disruption of larval body color, and lack of pigmentation in the adult eyes (Suppl. Fig3c). We did not detect any phenotypes from *vermilion* and *scarlet* mutants in larvae or adults, even though a previous study detected *vermilion* expression in developing wing tissue in this species (Beldade et al., 2005). We conclude that the ommochrome pathway is involved in coloring the larval body and adult eyes but likely not the wings of *B. anynana.*

### Deletion of melanin pathway genes affected scale morphology

To examine the contribution of melanin biosynthesis pathway products to the development of scale size and the cuticular microstructures of wing scales, we took SEM images of individual scales of different colors from male mutants and compared these to WT scales.

In *yellow* mutants, the windows of the black and brown scales were broadly covered by a supernumerary lamina (Fig.4a), which only rarely occurs in WT scales, with the exception of white scales, which naturally display parts of such lamina. White, gold, and beige scales showed no marked differences in these mutants. The crossribs flanking these closed windows in mutant scales were thinner, and the crossrib spacing was also significantly denser in black and brown scales. On the other hand, the distance between longitudinal ridges and overall scale size were unchanged (Fig.4b and Table).

**Figure 4.**
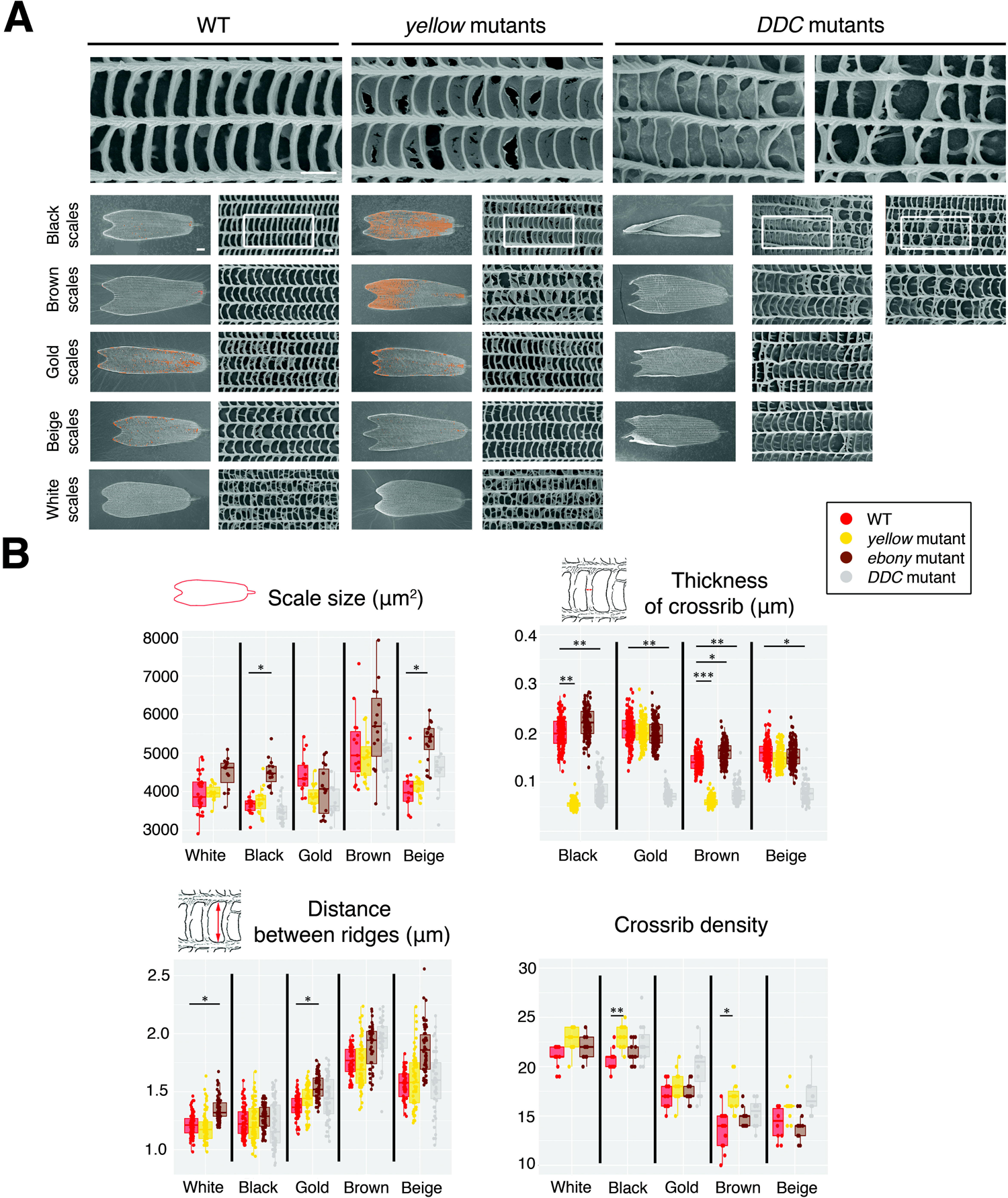
SEM images of individual scales from WT, *yellow* and *DDC* mutants and detailed morphological measurements from these as well as *ebony* mutants. Representative images of individual scales from WT, *yellow* mutants, and *DDC* mutants. In WT black, brown, and beige scales the windows, flanked by longitudinal ridges and crossribs, are open. In gold and beige scales, we can detect a thin lamina that partially covers a few of these windows. In white scales, the thin lamina partially covers many of the windows, and trabeculae are long and disorganized. In *yellow* mutants, there is a thin lamina that covers the windows of darker colored scales (black and brown) (the ectopic lamina was colored in orange in the SEM images). The morphology of the lighter colored scales (gold, beige and white) is unchanged. In *DDC* mutants, crossribs are more disorganized and sometimes fuse with neighboring crossribs, in addition trabeculae become sheet-like underneath the crossribs. These phenotypes were stronger in the darker colored scale (black and brown), and weaker in the lighter colored scales. Scale bars indicate 10 *μ* m in low magnification images of whole scales, and 1 *μ* m in high magnification images (top row). (B) Several measurements of individual scale features. We plot all measurements taken from five scales of the same type from three different individual but statistics used N=3, where the 5 measurements within an individual were averaged.

In *DDC* mutants, the arrangement of crossribs on all the color scales became disordered, and neighboring crossribs were often fused, resulting in larger windows. Most strikingly, sheet-like trabeculae, taller than the feet-like trabeculae of wild type scales, appeared beneath the crossribs. The width of each crossrib also became thinner (Fig.4 and table). On the other hand, scale size and distance between longitudinal ridges were unchanged. These phenotypes were stronger in darker colored scales (black), and weaker in lighter colored scales (gold and beige).

In *ebony* mutants, we did not detect dramatic change in morphology, but scale size was slightly larger in black and beige mutant scales, thickness of crossribs was slightly higher in brown mutant scales, and distance between longitudinal ridges was slightly larger in white and gold mutant scales, relative to WT scales (Fig4b). Most of these morphological changes had a small effect size, took place in scales where there was no significant affect on color, did not transform the morphology of one colored scale into that of another colored scale, and may be simply due to slight differences in genetic background between the six animals measured.

In summary, the color and the morphology of the darker scales in *yellow* mutants and the color and morphology of all the colored scales in *DDC* mutants were dramatically changed relative to WT scales. We also detected smaller changes in scale morphology in the *ebony* mutants, but these were small compared to the observed changes in color in these same mutants.

## Discussion

In this study, we generated mosaic mutants for five of the known melanin biosynthesis pathway genes and three of the ommochrome pathway genes using CRISPR/Cas9. This allowed us to identify which pigment pathway and which genes are needed for producing each of the colored scales on the wings of *B. anynana* and how these genes affect scale color and morphology.

### The role of melanin biosynthesis pathway genes in the development of scales and pigmentary colors.

The melanin pathway mutants displayed both expected color alterations, i.e., those observed in similar experiments in other insects and butterflies (True et al., 1999, Wittkopp et al., 2002, Arakane et al., 2009, Liu et al., 2010, Liu et al., 2014, Li et al., 2015, Liu et al., 2016, Perry et al., 2016, Zhang and Reed, 2016, Beldade and Peralta et al., 2017, Zhang et al. 2017), as well as previously undescribed color changes. A surprising discovery was that *aaNAT* is necessary for generating the complete skeleton of white eyespot center scales in *B. anynana,* which produce a white structural color (Monteiro et al. 2015). Interestingly, in *Bombyx, Oncopeltus,* and *Tribolium,* loss or down-regulation of *aaNAT* caused increased cuticle melanization (Dai et al., 2010, Zhan et al., 2010, Liu et al., 2016, Noh et al., 2016). We speculate that in those systems, *aaNAT* is co-expressed with *yellow* in the same cells, leading to the re-direction of the common dopamine precursor used by both enzymes to the exclusive use by *yellow* leading to the production of more dopamine-melanin and a darker color.

The color of the gold scales in *B. anynana* was affected by mutations in *ebony,* suggesting that NBAD sclerotin contributes to this color. However, we also observed that in *DDC* mutants, gold color persisted in the affected regions (Fig.3a). We speculate that pheomelanin may be an additional molecule contributing to this color in *B. anynana*. Pheomelanin is generated from either tyrosine or DOPA through different chemical reactions (Fig. 2) (Ito et al., 2008). Under the presence of cysteine, the precursor dopaquinone is used to produce pheomelanin. Despite the belief that invertebrates don’t produce pheomelanin, this product was recently detected in mollusks, grasshoppers, butterflies, bumblebees and wasps (Speiser et al., 2014, Galván et al., 2015, Jorge Garcia et al., 2016, Polidori et al., 2017) and may also be present in the gold wing scales of *B. anynana*.

Our experiments also clarified the role of *yellow* in the melanin biosynthetic pathway. In *B. anynana, yellow* mutants were still capable of generating brown color, visible throughout the wing but especially in the black eyespot area. According to Fig. 2, this color is likely to derive from dopamine-melanin. The *TH* and *DDC* mild mutants described above (Fig.3), by leading to whitish scales throughout, also point to dopamine-melanin as the only likely brown pigment on the wings. This indicates that *yellow* is not involved in producing dopamine-melanin from dopamine in *B. anynana,* and perhaps more broadly across insects, as previously hypothesized (Han et al., 2002). The function of *yellow* appears to be the same as that of *yellow-f* or *yellow-f2* in *Drosophila,* which can convert dopachrome into dihydroxyindole (DHI), but incapable of converting dopaminechrome into 5,6-dihydroxyindole-2-carboxylic acid (DHICA) (Han et al., 2002). Other, yet uncharacterized, yellow family genes might catalyze this latter reaction in *B. anynana* and in other insects.

The development of *B. anynana* wing scales appears to require both TH and DDC, two early component of the melanin pathway that severely also impacted scale sclerotization. Without *TH*, scale cells were unable to produce a cuticular skeleton. We could not distinguish, however, whether the absence of wing scales in severe *TH and DDC* mutant mosaic patches was caused by inhibition of wing scale development or the loss of developed scales upon emergence, due to the absence of a scale skeleton. Without adequate levels of either TH or DDC, scales developed a curly appearance and also had overall lower levels of pigmentation. These results indicate that expression of both genes is important in the development of the cuticular skeleton as well as in the development of melanin pigments.

Color on the wings of *B. anynana* appears to derive primarily from melanin pathway products where the different color patterns likely result from the different spatial, and perhaps also levels of expression of the different melanin pathway genes in each scale. By examining the affected colored regions in each mutant we inferred the spatial expression for each of these enzymes (Fig 5a). We propose that *TH* and *DDC* genes are expressed across the whole wing. *aaNAT* is expressed only in white scales of the eyespot center. *yellow* is expressed in both the background brown and black eyespot regions, where the differences in color between these two regions likely result from either more *yellow* and/or less *DDC* expression levels in the black scales. *ebony* is primarily expressed in both the background beige and gold eyespot regions. We further propose that the particular combination of enzymes expressed in each scale cell determines the direction and flux of the precursors of the branched melanin pathway (Fig.5a).

**Figure 5.**
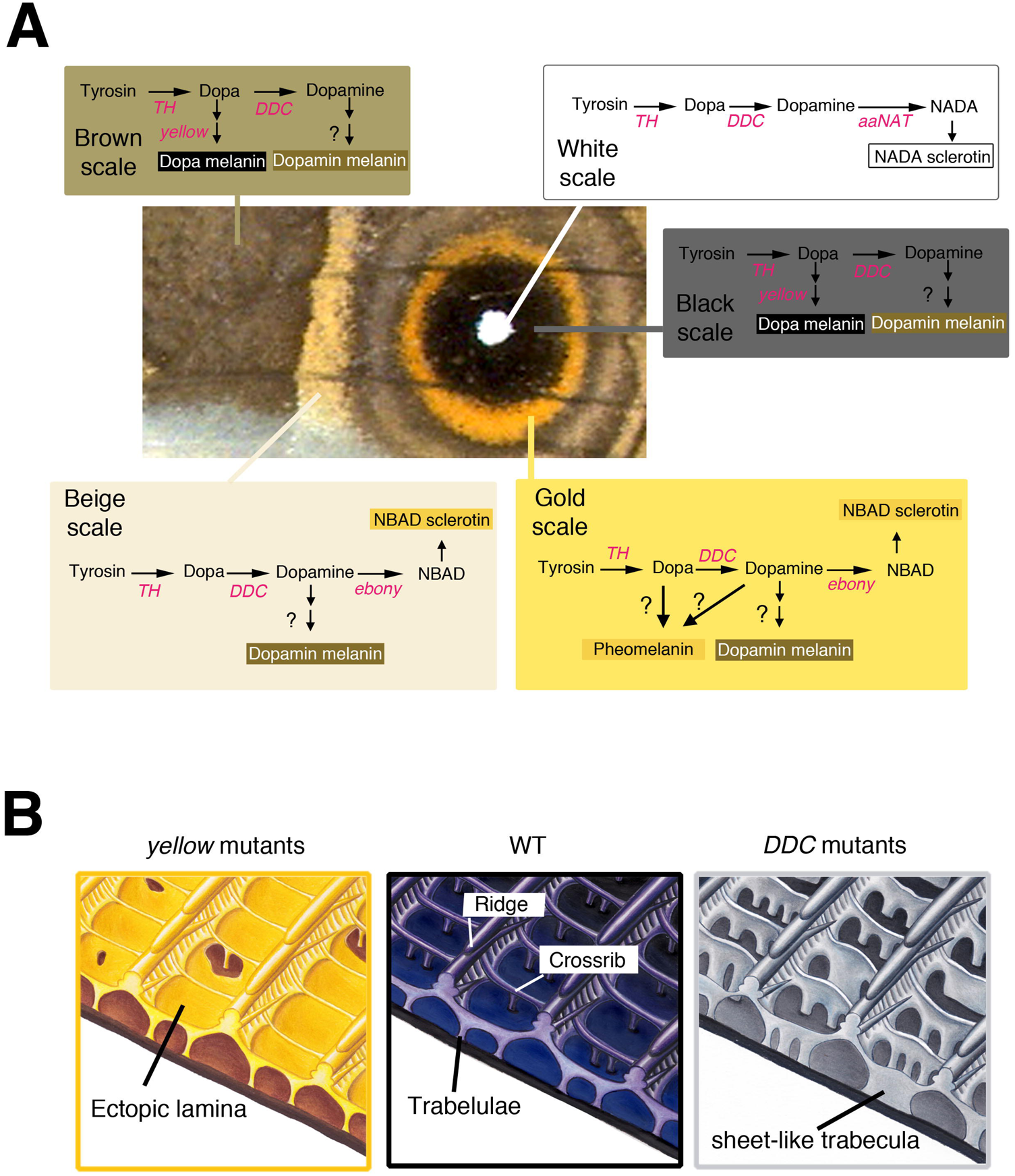
Cell-specific expression and function of melanin biosynthesis pathway genes in wing scales of *B. anynana* controls scale color and morphology. (A) Summary of the scale-cell specific expression of each of the examined melanin pathway genes, inferred from the loss of function mutants, on the wings of *B. anynana,* and how their combinatorial expression contributes to the production of different pigments in each scale cell. (B) Schematic diagram of the morphology of a black scale from WT, *yellow* and *DDC* mutants. In *yellow* mutants, black scales become yellow-brown, the normally open windows become covered by an ectopic cuticular lamina, and the adjacent crossribs became thinner, implicating the gene *yellow* in the creation of both dark pigmentation and in opening windows. In *DDC* mutants, the black scales become greyish, the trabeculae became sheet-like and taller, and the adjacent crossribs became thinner, implicating the gene *DDC* in the creation of dark pigmentation and short and pillar-like trabeculae (hand illustrations by Katerina Evangelou).

Previous wing color patterning work on this species, has primarily focused on the early molecular signals and transcription factors that set up the white center and the rings of color in an eyespot, in the late larvae and early pupal wing, several days before the expression of the pigment enzymes examined here (Brakefield et al. 1996, Brunetti et al. 2001, Monteiro et al. 2006, Saenko et al. 2011, Monteiro et al. 2013, Tong et al. 2014, Ozsu and Monteiro 2017, Özsu et al. 2017). Here, by uncovering the specific spatial distribution of five melanin pathway enzymes, we can now suggest a possible mapping between the expression of these early developmental genes and the enzymes at the end of the wing patterning process. In particular, *DDC* being expressed overall on the wing and *yellow* being expressed in darker colored scales correlates nicely with functional and quantitative *Distal-less* gene expression (Brakefield et al. 1996, Monteiro et al. 2013, Connahs et al. 2017), suggesting that *DDC* and *yellow* expression might be regulated by *Distal-less.* Any one of the 186 genes recently found differentially expressed in the central cells of an eyespot during the early pupal stage (Ozsu and Monteiro 2017) or at other time points (Monteiro et al. 2006, Özsu et al. 2017) could be up-regulating *aaNAT* expression specifically in these cells. Expression of *ebony* in the gold ring could be up-regulated by *engrailed* (Brunetti et al. 2001), but it is still unclear what could be regulating this gene in the beige transversal band. Finally *TH* (and *DDC*), might not be spatially regulated by any of the above pattern-associated transcription factors.

### Subcellular localization and functions of melanin pathway products during wing scale development

Butterfly wing scale development has been largely visualized via transmission electron microscope (TEM) images of fixed developing scale cells (Ghiradella et al., 1972, Greenstein 1972a,b, Ghiradella, 1989), but more recently fluorescent microscopy was also used (Dinwiddie et al., 2014). In TEM images of early scale development of *Ephestia kuehniella* moths, microfibril bundles, aligned with regular spacing, were observed on the surface of scale cells as the scale started to flatten (Overton, 1966). Greenstein proposed that the scale ridges of the Giant Silkmoth, *Hyalophora cecropia,* were likely to develop in between these microfibril bundles by buckling of the expanding extra-cellular cuticle (Greenstein, 1972b). Recently, Dinwiddie et al. (2014) showed that cuticle deposition along the future longitudinal ridges happened in between flanking longitudinal actin bundles inside the developing scale, and disruption of actin filament formation led to disorganized scales. Together, these studies indicate that scale ridge development likely depends on the interaction of actin filaments and cuticle. On the other hand, little is currently known about the molecular and genetic mechanism for cossrib and trabecular development.

The *yellow* and *DDC* mutants described here show how disruptions to the melanin synthesis pathway impact both scale color and morphology (Fig4). Even though interaction of dopa-melanin and dopamine-melanin, and other cuticule components have not been described so far, we infer that loss of dopa-melanin and dopamine-melanin are causing the *yellow* and *DDC* color and scale structural changes, respectively. We speculate that cuticle is first distributed at the periphery of the developing scale, including in the window regions, and the presence of dopa-melanin helps cuticle polymerize around the crossribs to create windows devoid of cuticle. Similarly, cuticle would be present initially in vertical walls below the cross-ribs and the presence of dopamine-melanin in these areas would help polymerize cuticle more tightly under and around each crossrib, to create short and thick pillars under the ridges, the trabecullae, as well as thicker crossribs (Fig.4). At later stages, the cytoplasm of the scale cells disappears leaving behind the cuticular skeleton (Ghiradella, 1989). Cells that don’t express *DDC* leave behind tall, sheet-like trabeculae, and cells that don’t express *yellow* leave behind unaggregated cuticle as a thin lamina covering the windows. We propose that the polymerization and interaction of sub-cellular compartmentalized dopa-melanin, and dopamine-melanin molecules with chitin or other cuticular proteins is important for constructing specific cuticular morphologies in the scale cell, especially involving cuticle aggregation around trabeculae and cross-ribs. The presence of parts of horizontal laminae, with multiple holes, is also visible predominantly in WT white scales, where *yellow* expression was inferred to be naturally absent. The extra chitin lamina in *yellow* mutant scales is reminiscent of the silver scales found in *Agraulis vanilla* (Dinwiddie et al., 2014) or in the scales of *P. palinurus,* in which the windows are completely filled by multiple layers of laminae (Vukusic et al., 2000). It will be interesting to explore how melanin pathway genes affect the morphology of the scales in these other species.

Mutations in melanin pathway genes also affected scale morphology in subtler ways, such as in altering the spacing between ridges and crossribs but these changes were not sufficient to explain why scales of different colors exhibit these different morphologies (Suppl. Fig. 1 and 2, Janssen et al., 2001). These results suggest that other factors, besides melanin pathway products, are regulating these subtler quantitative differences in scale morphology between scales of different colors.

Some of these factors might involve overall concentration of cuticle inside the cell and type of cuticular proteins expressed in each scale. Ohkawa et al., 2004 demonstrated that electrospun fibers formed by lower concentrations of chitosan, a derivate of chitin, were thinner than those formed at higher concentrations. It is possible, then, that eyespot-forming scales, showing a higher density of crossribs, might have higher concentration of cuticle building materials than beige or brown scales. Finally, the amount and/or the spatial distribution of the large number of cuticle proteins found in lepidopterans (Liang et al., 2010), which have been recently implicated in the joint regulation of larval body cuticle thickness and its coloration (Xiong et al. 2017), might also contribute to particular scale morphologies.

## Conclusion

In this study, by disrupting melanin and ommochrome pathway genes we demonstrated that most of the color observed on the wings of *B. anynana* is comprised of melanin biosynthesis pathway products, and showed how each of the colors is due to the expression of a specific sub-set of melanin pathway enzymes in each scale cell, that channel the flux of precursors in the pathway into specific end pigments. This work, thus, contributes to our understanding of the genetic mechanism that establish complex color patterns in butterfly wings. We also showed that melanin biosynthesis pathway products are acting both as pigments and as scaffolds to build the fine cuticular structures of a wing scale. These are the first molecules described, together with actin (Dinwiddie et al., 2014), that affect the microstructures on a wing scale. The genes examined here should perhaps also be investigated in other butterflies, in particular those that show complex morphologies (Saranathan et al. 2010) and iridescent structural colors (Ghiradella et al., 1972, Vukusic et al., 2000) as they may contribute to the construction of complex photonic materials. Uncovering the genetic basis of such finely sculpted biophotonic materials may eventually pave the way for a live-cell bioengineering process of fabricating these materials.

## Materials and Methods

### Butterfly husbandry

*B. anynana*, originally collected in Malawi, have been reared in the lab since 1988. Larvae were fed on young corn plans and adults on mashed banana. *B. anynana* were reared at 27°C and 60% humidity in a 12:12 light:dark cycle.

### sgRNA design

sgRNA target sequences were selected based on their GC content (around 60%), and number of mismatch sequences relative to other sequences in the genome (> 3 sites). In addition we picked target sequences that started with a guanidine for subsequent in vitro transcription by T7 RNA polymerase.

### sgRNA production

Template for in vivo transcription of sgRNA was made with PCR method described in Bassett et al., 2014. Forward primer contains T7 RNA polymerase binding site and sgRNA target site (GAAATTAATACGACTCACTATA**GNN19**GTTTTA GAGCTAGAAATAGC). Reverse primer contains the remainder of sgRNA sequence (AAAAGCACCGACTCGGTGCCACTTTTTCAAGTTGATAACGGACTAGCCTTAT TTTAACTTGCTATTTCTAGCTCTAAAAC). PCR was performed with Q5 High-Fidelity DNA Polymerase (NEB) in 100ul reaction volumes. After checking with gel electrophoresis, PCR product was purified with Gene JET PCR purification kit (Thermo Fisher).

In vitro transcription was performed with T7 RNA polymerase (NEB) using 500 ng of purified PCR product as a template for overnight. After removal of template DNA with DNase1 treatment, the RNA was purified with ethanol precipitation. The RNA was suspended to RNase-free water and stored at −80 □.

### Cas9 mRNA production

Plasmid pT3TS-nCas9n (Addgene) was linearized with Xba1 and purified by Phenol/Chloroform and ethanol precipitation. In vitro transcription of mRNA was performed with mMESSAGEmMACHINE T3 kit (Ambion) using 1ug of linearized plasmid as a template, and poly(A) tail was added by using Poly(A) Tailing Kit (Thermo Fisher). The RNA was purified with lithium-chloride precipitation, and then suspended to RNase-free water and stored at −80 □.

### Microinjection

Eggs were laid on the corn leafs for 30mins. We co-injected 0.5 μg/μl final concentration of gRNA and 0.5 μg/μl final concentration of Cas9 mRNA into embryos within 1h after egg laying. Eggs were sunk into PBS, and injection were performed inside the PBS. Food dye was added to the injection solution for visualization. Injected embryos were incubated at 27□ in PBS, and transferred on the silk on the next day, and further incubated at 27 □. After hatching, larvae were moved to corn leafs, and reared at 27 □ with a 12:12 h light:dark cycle and 60% relative humidity.

### Detection of indel mutations

Genomic DNA was extracted from a pool of about 5 injected embryos that did not hatch with SDS and Proteinase K method. About 250 bp of sequence spanning target sequence was amplified with PCRBIO Taq Mix Red (PCRBIOSYSTEMS), and optimized the PCR condition that do not show any smear, primer dimer and extra bands. Primers for those analyses are listed on Table. PCR products were purified with Gene JET PCR purification kit (Thermo Fisher).

Two hundred ng of PCR product were denatured and reanealed in 10x NEB2 buffer. One ul of T7 endonuclease □ (NEB) were added to the sample, and 1ul of mQ was added to a negative control. Immediately after incubation for 15 min at 37 □, all the reaction was run on the 3% agarose gel.

The PCR product which showed positive cleavage result from T7 endonuclease □ assay was subcloned into pGEM-Teasy vector (Promega) with TA cloning. For each target, we picked 8 colonies and extracted the plasmid with traditional alkali-SDS method, and PEG precipitation. Sequence analysis was performed with BigDye Terminator v3.1 Cycle Sequencing Kit (Thermo Fisher) and 3730xl DNA Analyzer (ABI)

### Acquisition of colored images

The images of butterfly wings and larvae were taken with a LEICA DMS1000 microscope camera. Color images of individual scales were taken after embedding scales in clove oil. This oil matches the refractive index of insect cuticle and allows pigmentary color to me measured precisely (Wasik et al, 2015). In particular, this oil treatment removes any contribution of structural colors that might result from light bouncing and constructively interfering when crossing finely patterned materials with different refractive indexes, such as cuticle and air. First we put one drop of clove oil on a glass slide. Wing scales were taken from a mutated site (from the area depicted in Fig. 3B) by using a fine tungsten needle, and were dipped into the clove oil. Once enough scales were transferred, a cover glass was placed on top of the oil and the scales. Additional clove oil was supplied from the edge of cover glass to fill the space and this preparation was sealed with nail polish. Images of wing scales were taken via light transmission using an Axioskop2 mot plus (Carl Zeiss) microscope, and Axiocam ERc5s (Carl Zeiss).

### Color measurements

Transmission images taken by above method were not entirely consistent for brightness, so images were first edited to make the background brightness similar across all images using the DigitalColor Meter, which is a pre-installed Macintosh app. The same relative region in a scale (in the middle) was selected with a rectangular marquee of a constant size, and the color was averaged inside the selected area using the “average” tool in Adobe Photoshop (CS3). For quantitative assessment of color, we conducted L*a*b color space analysis, which is a way of mathematically describing color in three-dimensions with 3 parameters. The L value (0 to 100) indicates how bright the color is with higher values indicating a brighter/lighter color, the a value (-60 to 60) indicates where the color is located between red (negative values) and green (positive values) opponent colors, and the b value (-60 to 60) indicates where the color is located in between blue (negative) and yellow (positive) opponent colors. The L*a*b color space is used in industry and also in biological color assessment. Measurements were conducted using the DigitalColor Meter. In this study we plotted only the L and b values since the a value (red-green) was similar among all measured scales. Color measurements were averaged across 5 scales of the same color from the same individuals, and three different male and female individuals were measured for statistical purposes. ANOVAs were used to test for significant differences in mean color values between the sexes and between WT males and each of the male melanin mutants using IBM-SPSS Statistical version 21 software.

### Scale measurements

We compared several traits between males and female scales such as area, length, width, average distance between longitudinal-ribs, density of cross-ribs along a standard length, and width of each crossrib, in both cover and ground scales of different colors. For crossrib thickness, distance between crossrib we we took the average of 10 measurements in a different area of the scale, and for distance between ridges we took the average of 3 measurements in a different area of the scale, and for scale area size, scale width, scale length, and scale density we took the average of 1 measurements in a different area of the scale, and then the average of 5 individual scales of the same color from the same individual. Our sample size, for statistical purposes, was = 3, i.e., the measurements from three distinct individual animals of each sex. We measured each trait using Image J software. ANOVAs were used to test for significant differences in scale measurements between the sexes and between WT males and each of the male melanin mutants using IBM-SPSS Statistical version 21 software.

### Scanning Electron Microscopy (SEM) and sample preparation

Small pieces of wing (5 mm × 5mm) were cut under a dissecting microscope. Fifty percent of Ethanol/mQ water was dropped onto the wing pieces, and then the pieces were dipped into liquid nitrogen for 5 mins. The wing pieces were taken out from liquid nitrogen, and evaporated at room temperature. The pieces of wing or individual scales were moved using a sharpened tungsten needle onto a double sticky carbon tape and pressed gently onto the metal SEM stub. After enough desiccation (overnight), platinum conductive coating was performed during 60 seconds at 20 mA using an auto Fine Coaters JFC-1600. Images were taken using a JSM-6701F, JEOL scanning electron microscope.

---

**Supplemental Figure 1.**
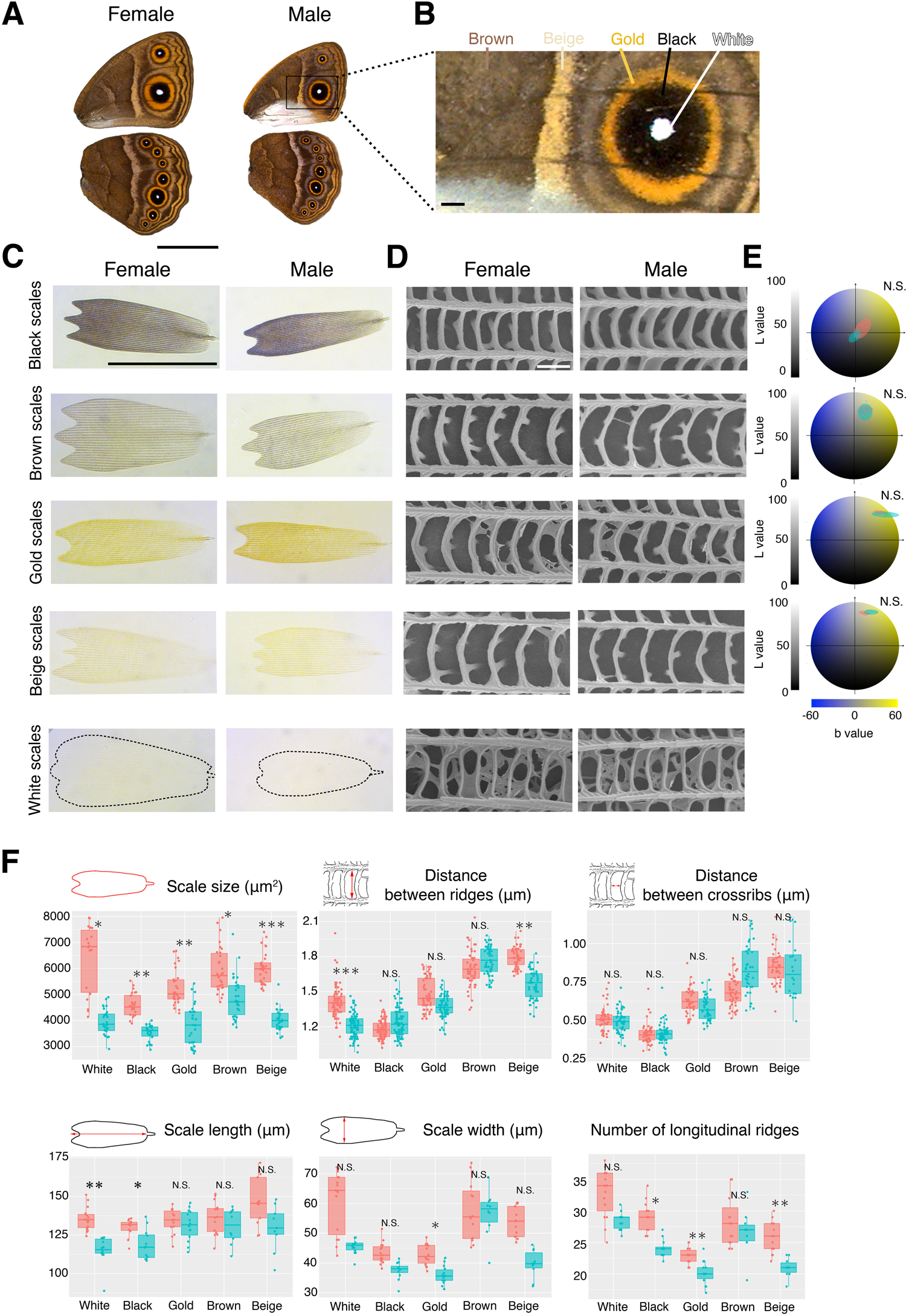
Males and females display differences is size and shape, but not color of cover scales. (A) Ventral surface of female (left) and male wings. Scale bar indicates 1 cm. (B) Colored areas that were sampled for individual scale measurements. Scale bar indicates 1 mm. (C) Transmission images of individual cover scales from males and females. Scale bar indicates 100 *μ* m. (D) Highly magnified SEM images of wing scales. Scale bar indicates 1 *μ m.* E) Color assessment of each scale using L*a*b color space. Red ellipses represent female data, and blue male data. There was no significant difference in scale color between the sexes. (F) Measurements of scale size, distance between longitudinal ridges, distance between crossribs, scale length, and total number of longitudinal ridges. We plot measurements from five scales from three different individual but statistics used N=3, where the five measurements within an individual were averaged. Females have consistently larger scales than males, for several other traits female dimensions are also larger than males, but for most traits sexes are similar.

**Supplemental Figure 2.**
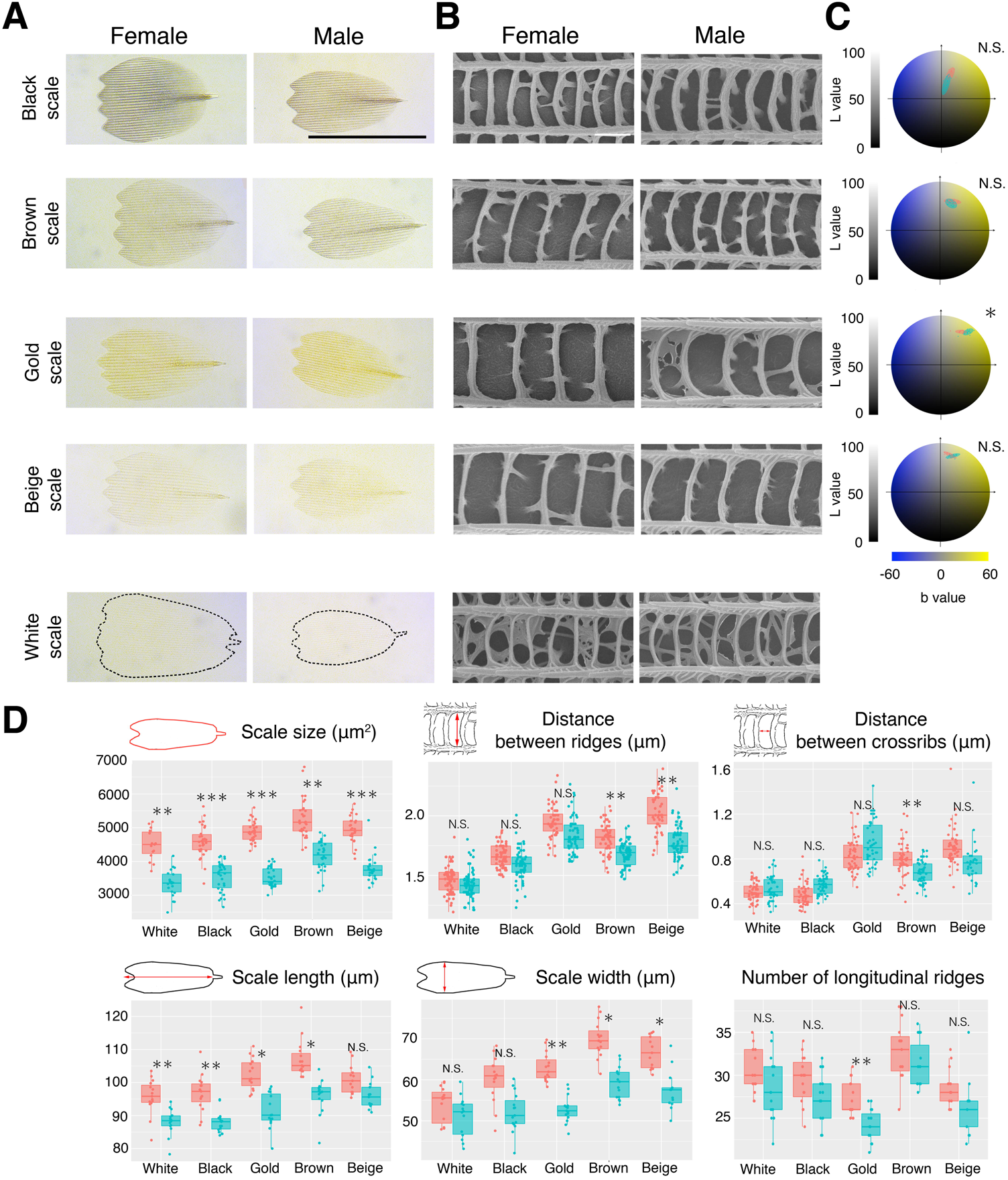
Males and females display differences is size, shape, and color of ground scales. (A) Transmission images of individual ground scales from males and females. Scale bar indicates 100 *μ* m. (B) Highly magnified SEM images of wing scales. Scale bar indicates 1 *μ* m. (C) Color assessment of each scale using L*a*b color space. Red ellipses represent female data, and blue represent male data. Differences in color are found in gold scales alone. (D) Measurements for scale size, distance between longitudinal ridges, distance between crossribs, scale length, and total number of longitudinal ridges (plotted as is Suppl Fig.1). Scale size is significantly larger in females, due to a different combination of variation in scale length and scale width (for different colored scales). Number of ridges, distance between longitudinal ridges, and distance between crossribs is larger for females for only a few of the colored scales.

**Supplemental Figure 3.**
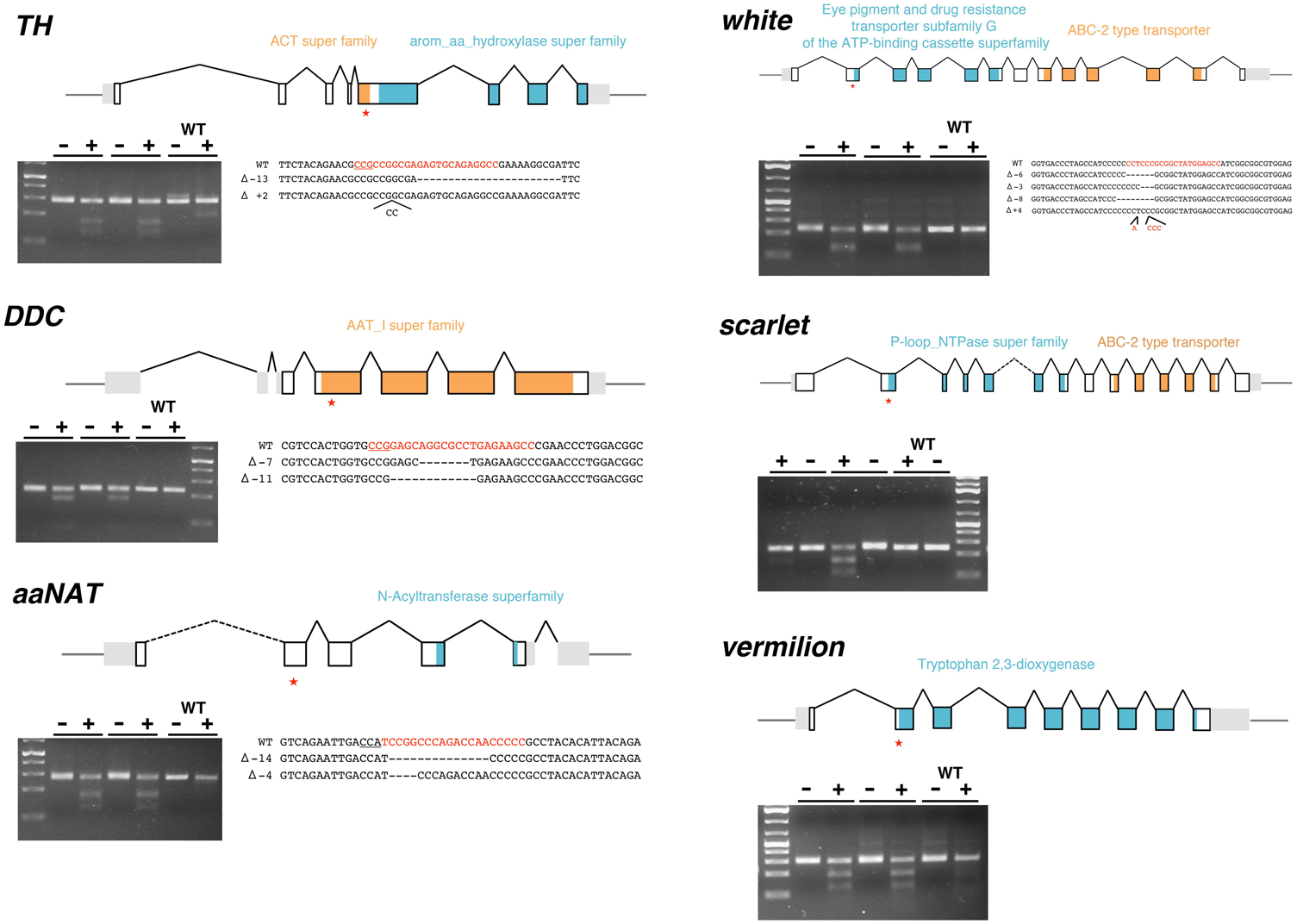
T7 endonuclease I assay and sequence analysis for CRISPR-Cas9 mutations. Schematic representation of gene structure of target genes, white boxes indicate exons, black v-shaped lines connecting boxes indicate introns, and gray boxes indicate untranslated regions. Colored regions inside exons indicate functional domains. Each functional domain was annotated using a conserved domain search at NCBI. Asterisks indicate the CRISPR target region. Gel shows the result of a T7 endonuclease assay performed on embryos after injection of guide and Cas9. Minus lanes indicate the no T7 enzyme reaction (negative control). Plus lanes indicate the reaction with the enzyme. Expected size of digested bands was observed only from the lane having T7 enzyme. Sanger sequence results indicate an indel mutation was generated around target site.

**Supplemental Figure 4.**
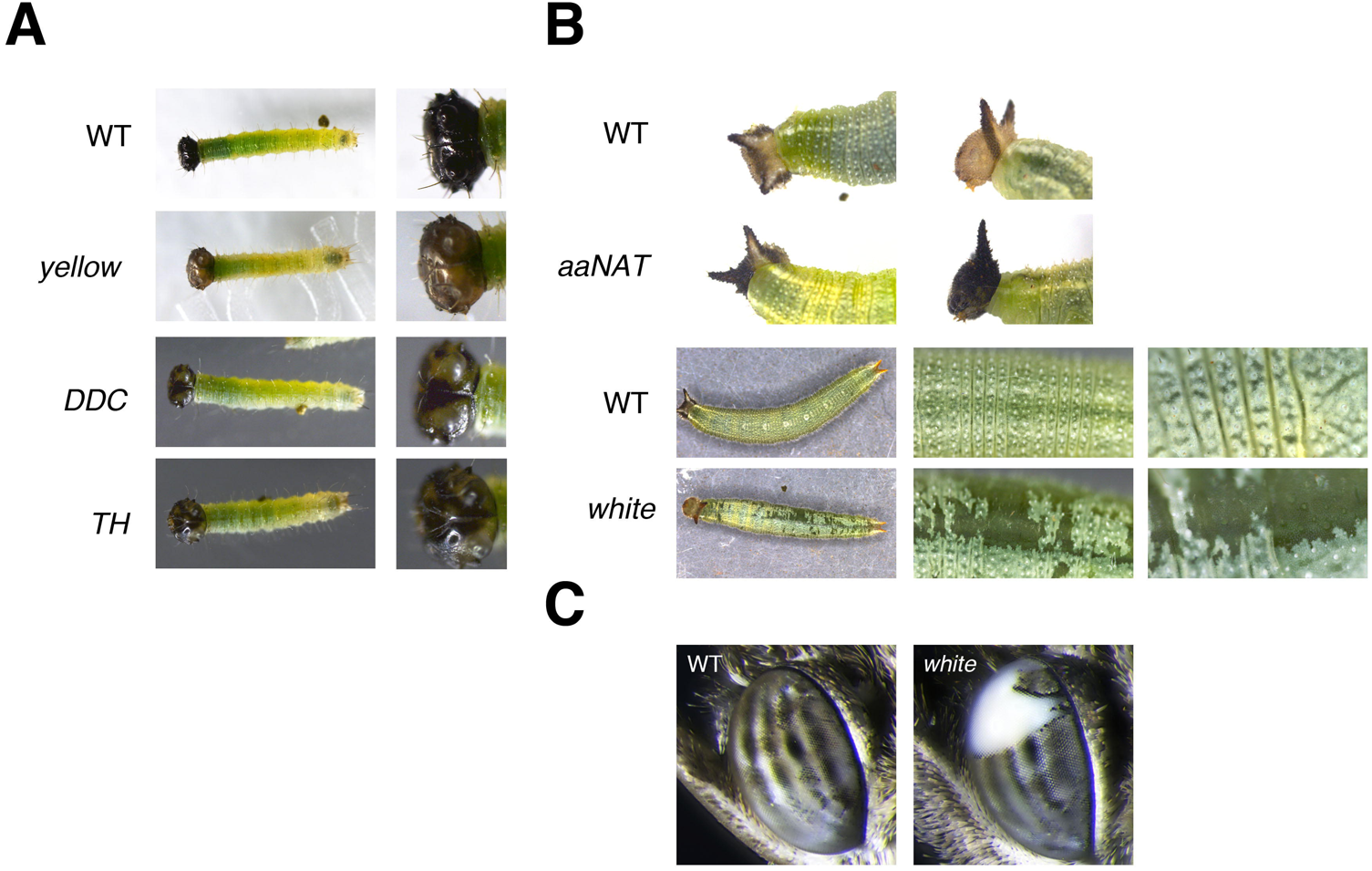
Melanin and ommochrome genes phenotype on larvae. (A) First and second instar larvae head capsule is usually completely black. In *yellow* mutants, partial head showed a brown color. In *TH* and *DDC* mutants, partial head lost pigment and became transparent. (B) Third instar larvae head capsule is usually brown. In *aaNAT* mutants, the head capsule became completely black. Third instar larvae body is usually whitish green. In *white* mutants, the larval body lost the whitish green color and became transparent. (C) Adult eye is usually colored in alternating black and white stripes. In *white* mutants, parts of the eye became completely white.

**Supplemental Figure 5.**
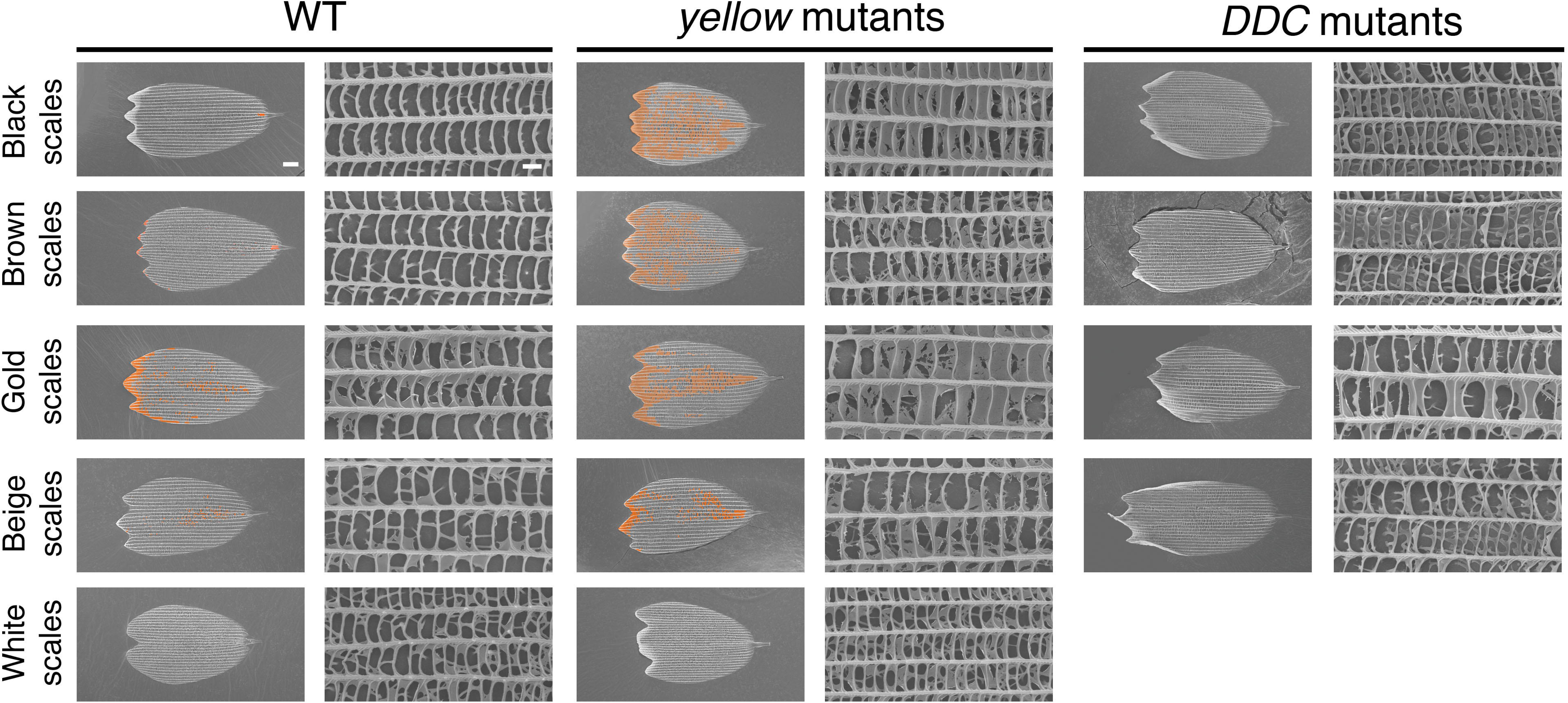
SEM images of individual ground scales from *yellow* and *DDC* mutants. Representative images of individual scales from WT, *yellow* mutants, and *DDC* mutants. Phenotypes were similar to those observed in cover scales. The ectopic laminae (colored in orange) and tall trabeculae in *yellow* and *DDC* mutants, respectively, were more extreme in the darker colored scales. Scale indicates 10 μ m in low magnification, and 1 μm in high magnification.

